# Determining salmon provenance with automated otolith reading

**DOI:** 10.1101/2021.10.14.464436

**Authors:** Chandler E Kemp, Susan K Doherty

## Abstract

Synthetic otolith marks are used at hundreds of hatcheries throughout the Pacific Rim to record the release location of salmon. Each year, human readers examine tens of thousands of otolith samples to identify the marks in salmon that are caught. The data inform dynamic management practices that maximize allowable catch while preserving populations, and guide hatchery investments. However, the method is limited by the time required to process otoliths, the inability to distinguish between wild and un-marked hatchery fish, and in some cases classification processes are limited by the subjective decisions of human readers. Automated otolith reading using computer vision has the potential to improve on all three of these limitations. Our work advances the field of automated otolith reading through a novel otolith classification algorithm that uses two neural networks trained with an adversarial algorithm to achieve 93% classification accuracy between four hatchery marks and unmarked otoliths. The algorithm relies on hemisection images of the otolith exclusively: no additional biological data are needed. Our work demonstrates a novel technique with modest training requirements that achieves unprecedented accuracy. The method can be easily adopted in existing otolith labs, scaled to accommodate additional marks, and does not require tracking additional information about the fish that the otolith was retrieved from.

**Author summary:** Many fish harvested in commercial fisheries have a bone-like structure in their inner ear called an otolith. Otoliths are useful because as they grow changes in the fish’s environment create unique patterns that can be interpreted years later—similar to the way tree rings can be used to determine a tree’s age and rate of growth in different seasons. Hatcheries use this phenomenon to create unique patterns in the otoliths of fish they release that identify their origin. Trained otolith “readers” examine tens of thousands of otolith samples as part of commercial fishery management every year. The data help resource managers protect returns to wild streams and help hatcheries track the survival rates of their releases. However, human reading is time consuming, limited in the types of patterns that can be identified and can be subjective. Applying modern machine learning techniques to otolith reading has the potential to improve on all of these challenges. We developed a computational method that identifies hatchery marks in digital images of salmon otoliths without human input. In a test with otoliths containing one of four hatchery marks and unmarked otoliths, the method accurately classified 93% of otoliths. The method can be adopted in otolith labs and can be further improved by expanding the sample size used to train and test the method.

## 1 Introduction

Sustainable salmon fisheries require complex and dynamic management policy, especially in the presence of climate change, habitat degradation and mixed wild and hatchery spawned fish [1]. Methods for quickly and accurately identifying salmon origins allow fishers to harvest hatchery salmon without threatening populations returning to wild streams [2], [3]. In addition, hatcheries benefit from identifying the release sites of fish to determine survival rates at different facilities. Otolith marking and reading has emerged as a popular tool for rapidly determining salmon origin in fisheries throughout the Pacific Rim [4].

Otoliths are a calcium carbonate structure found inside salmon ears that grow throughout their life. Changes in temperature, salinity or other environmental factors create more opaque or transparent layers within the otolith. When the otolith is extracted from a fish, sanded to a hemi-section and viewed under a microscope, the dark and light patterns can be “read” to learn about the salmon’s history, similar to the way tree rings can be counted to learn a tree’s age.

Hatcheries manipulate the temperature of water that fish are reared in to create unique patterns on the young fish’s otoliths. Since hatcheries release millions of fry at a time but only a few percent of the fry return to the fishing grounds as adults, the ability to mark all of the hatchlings simultaneously and economically makes otolith marking a popular tool for hatcheries. Figure 1 shows the location of hatcheries throughout the Pacific Rim that report their otolith marking patterns to the North Pacific Anadromous Fish Commission and the number of distinct marks expected to return to each hatchery in 2020 (see Supporting Information section S4 for additional details on the development of this figure) [5]. With billions of fish marked every year and millions returning to multiple hatcheries and wild streams, developing sufficiently distinct marks for humans to identify quickly is challenging.

**Figure 1.**
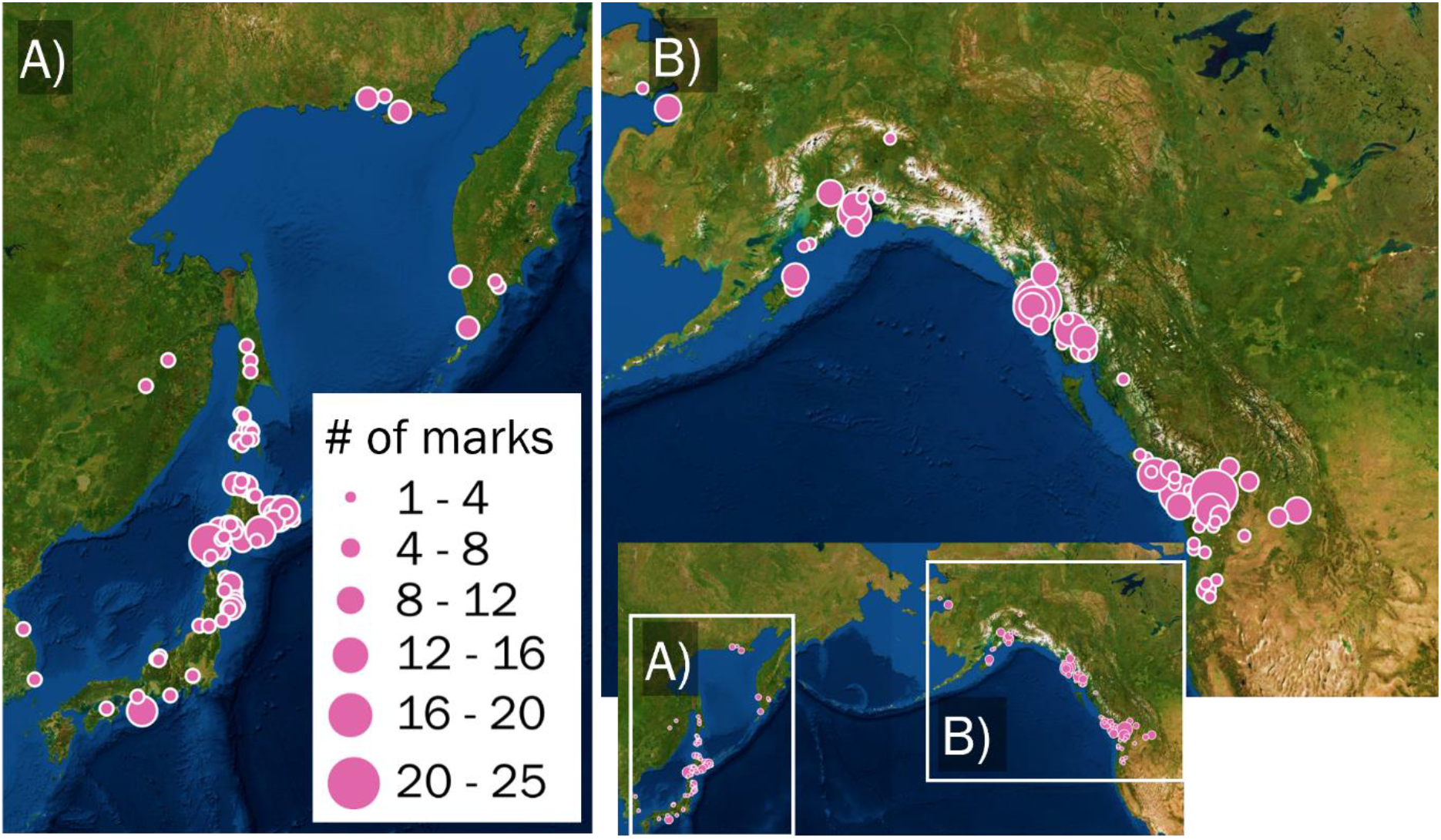
Location of hatcheries using otolith marking to identify salmon origins in the Pacific Rim including Japanese, Russian, American and Canadian locations. The pink circles indicate the location of hatcheries and the diameter of the circles indicates the number of unique marks returning to each site in 2020. In total, over five billion marked fry were released in years expected to return in 2020.

Every year, tens of thousands of otoliths are extracted from harvested salmon, cleaned, sanded, glued to a microscope slide and evaluated by human readers [6]. Experienced readers recognize the patterns developed by hatcheries and use them to determine which hatchery reared the fish. In other fisheries, otoliths are also used to estimate fish age or movement [4], [7]. The information collected from otoliths empowers fishery management to ensure adequate returns to wild streams and enables hatcheries to accurately assess the survival rate of their fish. However, management decisions are limited by the time required to process otoliths, the inability to distinguish between fish reared in different wild streams, and in some cases classification processes are limited by the subjective decisions of human readers. Automated classification schemes may help to address all three of these limitations.

Automated otolith reading methods have been studied almost as long as otoliths have been used for fishery management. Early methods used a one dimensional sample of an otolith image and applied pattern recognition algorithms to determine the number of rings and estimate fish age [8]. Between 1990 and 2000 several algorithms were published that incorporated increasing amounts of otolith structural knowledge into aging and classification schemes. Troadec [8] introduced a growth function before applying spectral analysis to account for the declining radial growth rate as fish age. With the growth function, Troadec achieved age estimations with a standard error of 1.6 and 3.8 days in hatchery and in situ samples for fish ranging from 14 to 35 days of age. Rodin et al. recognized that dark otolith bands are typically parallel and used this observation to reconstruct missing parts of bands [9]. Troadec et al. expanded on this concept by applying a ‘deformable template’ that parallels the circumference of the otolith to digital images. Age rings were identified based on the image brightness along the template [10].

Volk et al published one of the earliest otolith mark reading algorithms [11]. In the experiment, thermal marks were imposed on Chinook salmon otoliths. Hemi-sections of the otoliths were imaged and an algorithm for detecting the bands based on optical density gradient was tested. The algorithm failed to correctly locate the bands in 90% of otoliths. Subsequently, thin sections were produced from the sample, and the algorithm was tested again. In this case, bands were correctly located in 80% of otoliths, and the band locations were entered by hand in the remaining cases. Once the band locations had been corrected, the algorithm achieved perfect classification. However, the preparation of thin sections was significantly more time consuming than the preparation of hemi-sections; the method required human input; and the automated classification method was more time intensive than manual classification.

Since 2000, research focus has shifted away from algorithms that count rings directly toward more computationally intensive machine learning methods. In particular, several papers apply neural networks to otolith aging problems. Morison and Robertson developed a neural network for aging fish that included both otolith image data and biological data such as length and weight [12]. The neural network ages were compared with standard manual age methodologies [13]. The results showed that using the otolith image data only, neural network approaches had an average percent error in age estimate of approximately 20%. Including the biological data reduced that error to 5-10%. Fablet et al. compare results of fish aging based on neural networks and support vector machines (SVM) [14]. The neural network achieved approximately 82% agreement with an expert reader while the SVM approach using the same training set and similar features achieved an 88% agreement.

Deep convolutional neural networks with transfer learning have emerged as effective tools for classifying images with small training data sets [15]–[18]. We used transfer learning to adapt the VGG19, Inception and Xception networks to otolith mark classification, but found that using a pair of shallow networks to identify marked images and classify the marks achieved better classification accuracy with the available training set. The final algorithm presented here uses one network to select adversarial samples to fine-tune training of the second network, similar to the approach often used to train generative networks [19].

We present a novel otolith classification algorithm that uses two shallow neural networks trained with an adversarial algorithm to achieve 93% classification accuracy between four hatchery marks and unmarked otoliths in a test set with 100 images. No information about the fish is required by the algorithm beyond hemisection images of the otolith itself. This method advances the field by demonstrating a novel technique with modest training requirements that achieves unprecedented accuracy. The method is promising because it can be easily adopted in existing otolith labs, can be scaled to accommodate additional marks, and does not require tracking additional information about the fish that the otolith was retrieved from. We compare results with deep convolutional neural networks often used for image recognition and find the shallow networks achieve better sample classification accuracy given the small training database available for this work. Future work will explore the ability to improve classification accuracy by expanding the training set, the impact of lower quality otolith images on classification accuracy, and the persistence of classification accuracy as fish age.

## 2 Methods

We developed training and test sets from 250 otoliths collected from salmon reared in Southeast Alaska. Each otolith was sanded to a hemisection then imaged with 400x magnification. Hatchery marks are named using the notation illustrated in Figure 2. Individual dark patterns are referred to as “rings”. A collection of rings with consistent spacing create a “band.” Each mark can consist of one or more bands. The spacing between rings may be different in different bands, in which case the band with narrow spacing is noted with an “n”. Spacing between rings varies between otoliths with the same mark and between hatcheries. Therefore, the “n” notation is only used when two bands within the same mark have different spacing. In this work, we considered 50 otoliths each from five different marks: 3,5H10; 1,6H; 6,2H; 4n,2n,2H; and unmarked. Examples of each mark are provided in Figure 3.c-g. The training set included thirty images from each class. The remaining 20 images in each class were reserved for testing.

**Figure 2.**
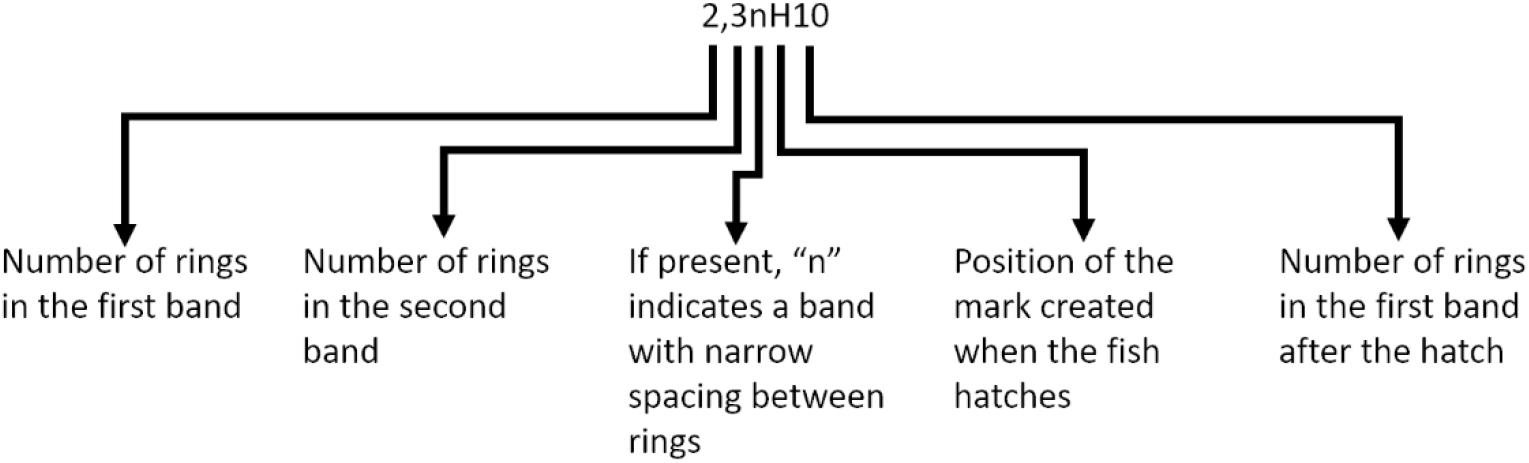
Illustration of otolith mark naming convention. The “n” notation is only used when ring spacing in one or more bands within a mark is intentionally made tighter than other bands in the same mark.

**Figure 3:**
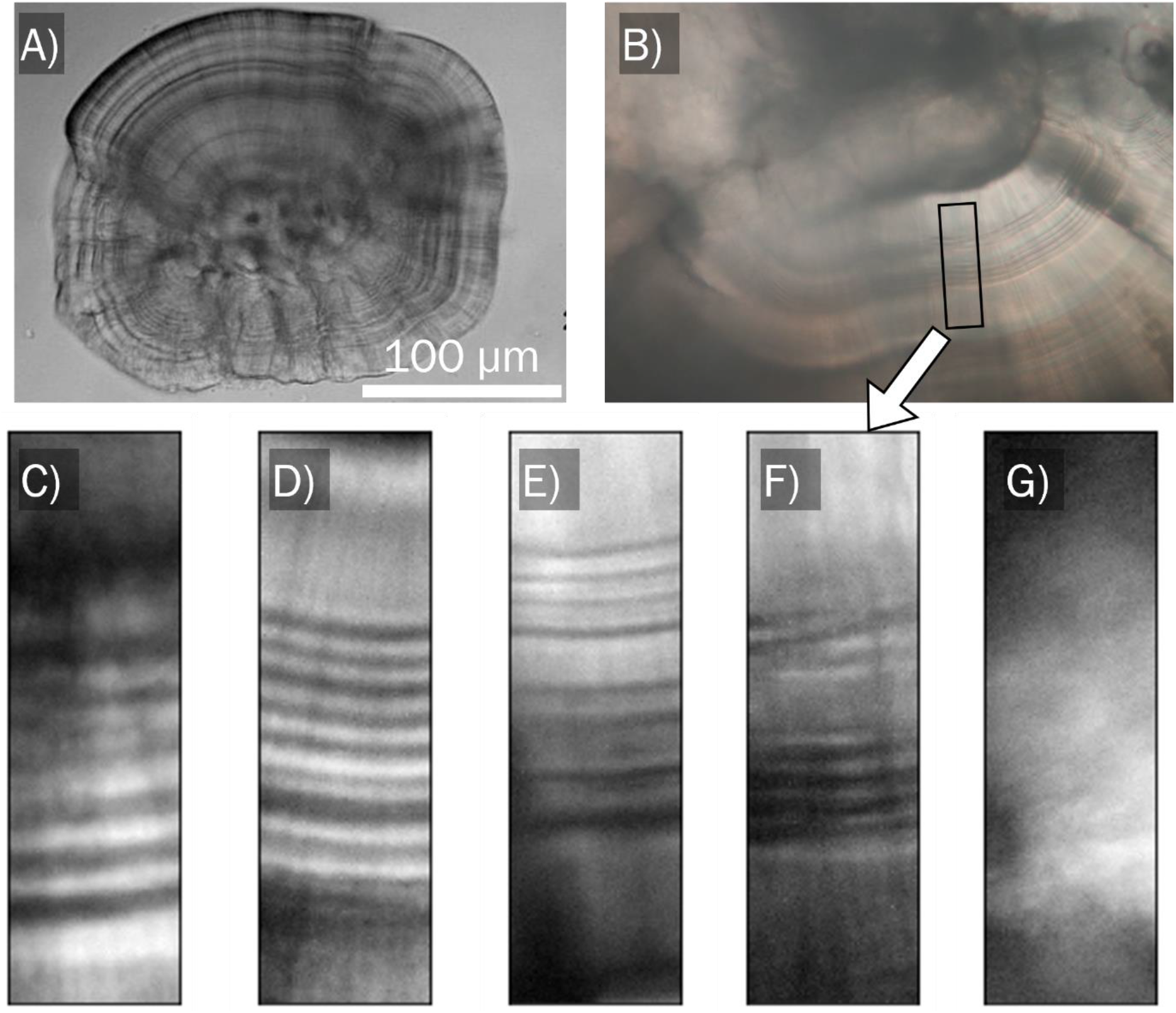
Marked otolith samples. A.) Whole otolith from a chum salmon before hatching. B.) An otolith hemi-section with a 3,5H10 mark viewed under a microscope. C.) 1,6H mark D.) 6,2H mark E.) 4n,2n,2H mark F.) 3,5H10 mark extracted from Figure 3.B. The post-hatch band is excluded from the sample. G.) Unmarked sample at the same magnification as samples C-F.

Each image has a resolution of 6000×4000 pixels. 240×800 pixel samples centered on the mark in each image were extracted using human input to specify the location of the mark to develop a set of training samples. The sample height (800 pixels) ensures all of the pre-hatch bands from every mark can be contained in one sample. The sample width (240 pixels) was selected to be narrow enough that the curvature of the rings across the sample was small compared to the width of the rings, but still wide enough that noise in the image could be overcome by averaging across the width of the sample. We developed software to support the extraction process publicly available through our Github repository [20]. Approximately thirty samples were extracted from each image, for a total of 800-900 training samples in each marked class. 120 unmarked samples were drawn at random from unmarked otoliths. Figure 3 illustrates the scale of a complete otolith in comparison to an otolith image used in this work and the samples extracted to highlight the marks. Each empirical sample was then used to generate 20 synthetic samples based on reflections, rotations of ±10 degrees and translations of ±2% of the sample height.

Otolith marks have three helpful characteristics that allowed us to reduce the dimensionality of the samples. Otolith marks are:

1. Approximately one dimensional: the signal does not vary across the width of a sample
2. Smooth: rings remain visible after smoothing and downscaling the images
3. Intensity and intensity gradient invariant: image brightness and monotonic changes in intensity across the height of a sample do not affect classification

To take advantage of these properties, the samples were averaged across their width, convoluted with a second derivative kernel, smoothed and then down sampled to an array of 151 numbers. A visualization of the preprocessing steps is provided in Figure 4.A-C. The processed samples were used to train a neural network with two convolution layers as shown in Figure 4.D. The resulting classification network is named the “Otolith Sample Classifier” (OSC).

**Figure 4 .).**
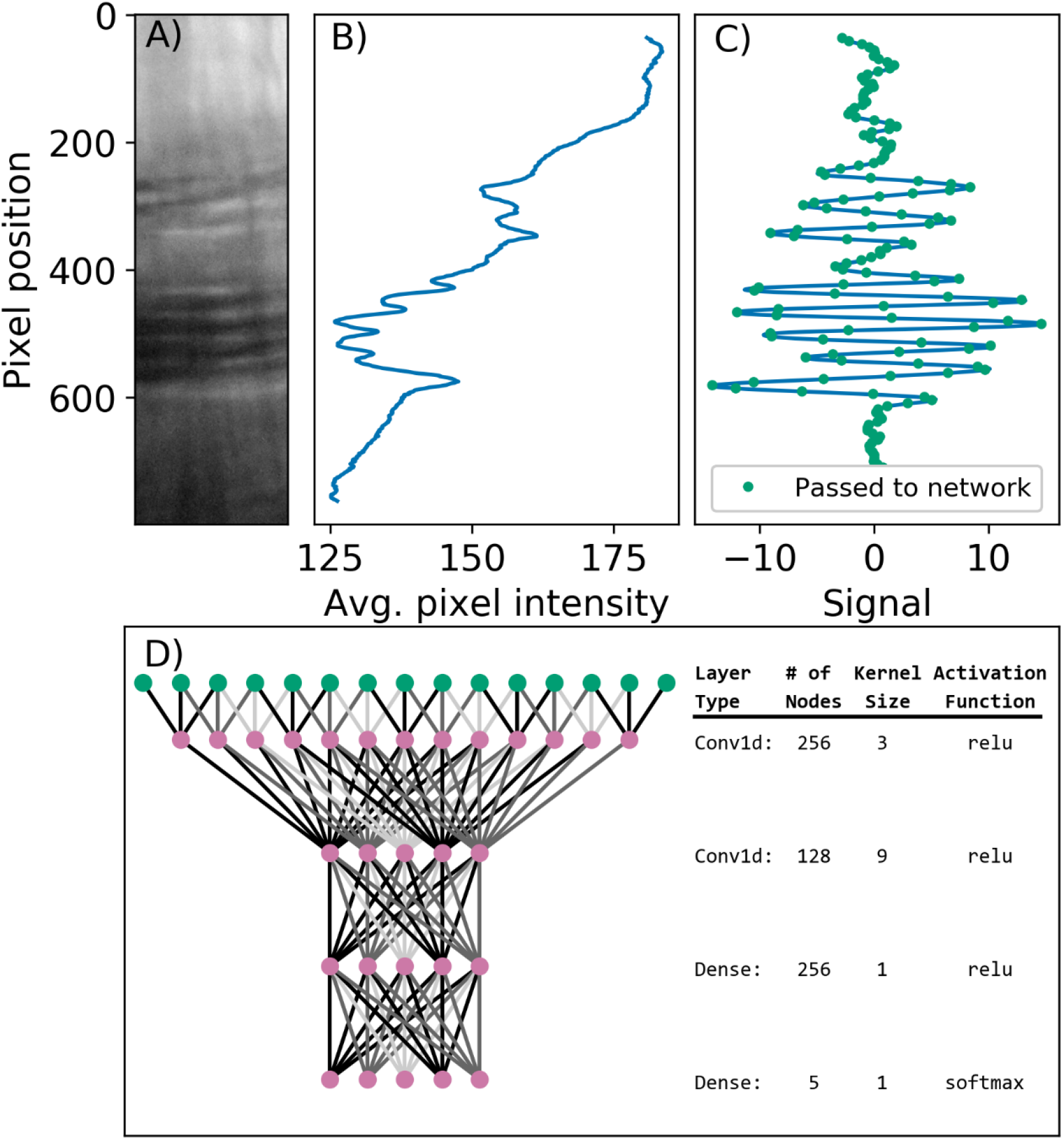
Illustration of the sample classification algorithm. A.) Example sample of the prehatch bands from a 3,5H10 mark. B.) Pixel intensity averaged across the x-axis of Figure 4.A on a scale of 0-255. C.) Signal developed after convolving the pixel intensity from panel B with a second derivative smoothing kernel. The data are down sampled to include every fifth pixel, as indicated by the green dots. D.) Visualization of the neural network used to classify samples.

The classification network described above provides a valuable tool for classifying samples, but it falls short of full image classification. Naively applying the network to samples drawn from an image fails to accurately distinguish between marked and unmarked images because the marks occupy a small fraction of the total image: iterating across an image with a step size of 120 pixels results in 1600 samples being drawn, of which approximately 30 will contain the mark in a marked image. In an unmarked image, a network that accurately identified 99% of samples as unmarked would still classify sixteen samples as marked. Indeed, when attempting to classify whole images using the classification algorithm described above, the network fails to identify unmarked images.

We overcame this limitation by training a second network to identify unmarked images. The second network requires the same type of input but returns a binary result: marked or unmarked. We refer to this second network as the “binary network.” The binary network is preliminarily trained on the same set of samples used to train the classification network. The classification algorithm is then used to automatically select samples from unmarked images in the training set that are identified as marked. We fine-tune the top layer of the binary network to discriminate between true marks and these adversarial samples. Fine-tuning is widely employed to use the features developed from large training sets to classify related but different samples with smaller training sets [21]–[23]. We adopted the method here by training features using the full training set and fine-tuning classification with a smaller training set of difficult to classify samples. We name this approach the “Adversarial Otolith Classifier” (AOC). All software to build and train the networks is available through Github [20]. The image database and trained networks are available through the SEANOE data repository [24].

### 2.1 Ethics statement

Ethical review and approval for use of animal tissue in this study was not required because all images were developed using otoliths previously collected in the normal operation of a commercial fishery.

## 3 Results

We report the accuracy of our algorithms in terms of the *sample* accuracy and the *image* accuracy. The sample accuracy defines the fraction of samples correctly classified by an algorithm. The image accuracy reports the fraction of images correctly classified by an algorithm. To evaluate image accuracy, an algorithm must accept a complete 4000×6000 pixel image as an input and return the most probable class for the otolith. The algorithm must include a process for identifying samples in the image that contain a mark, classify each of those samples and then combine the results from all samples to produce a single image classification.

We report results of both cross-validation and test-set experiments. In cross validation, a small fraction of the training data set is withheld for determining the accuracy of the model iteratively. This is called the “validation set.” In this case, we withhold 10% of the training set (15 images) for validation. We then iteratively rerun the validation experiment withholding a different set of 15 images until all of the images in the training set have been used for validation exactly once. Since this requires 10 iterations, the process is termed 10-fold cross validation. Cross-validation was performed many times in this work to test a variety of algorithms and tune settings before finally arriving at the algorithm presented here.

The test set was only exposed to the algorithm once it was finalized. Withholding the test set until the end of the experiment prevents the accuracy from being overstated due to iteratively modifying the algorithm to perform well on the validation set specifically.

Figure 5 shows the cross validation sample classification accuracy of the algorithm described above in comparison to the Xception, Inception and VGG19 networks commonly used for image classification [17], [18] (for further details describing how these networks were adapted for use in this problem, see SI section S2.3). The boxplots indicate the median and range of accuracies observed in ten-fold cross validation. The otolith-specific classification algorithm achieves a higher classification accuracy than the standard networks. The otolith-specific algorithm outperforms the other networks because it uses subject knowledge to reduce the dimension of the input data before training the network. The dimension reduction allows the classification network to be trained more effectively with a small training data set.

**Figure 5.**
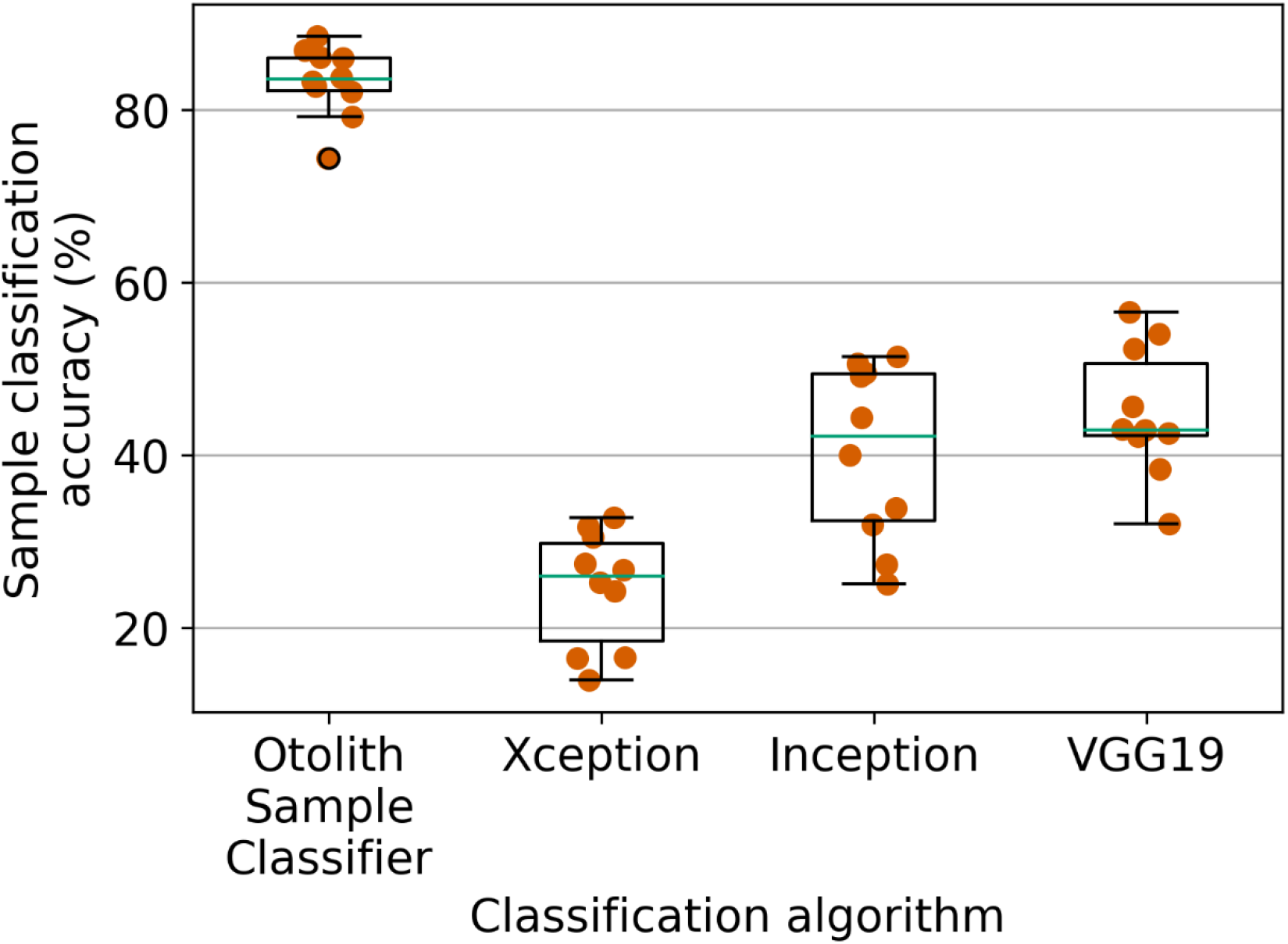
Sample classification accuracy of several tested networks.

A larger training set may allow the deep VGG19, Xception and Inception networks to be trained more effectively. However, since the signal in the otolith mark is essentially one-dimensional, traditional image classification networks should not be expected to perform as well as the customized OSC network. The two dimensional inputs of traditional image classification networks are quickly overfit to the sample size available here (see Figure S4 in SI).

Full image classification allows for some improvement over the sample classification accuracy because numerous marked samples can be extracted from every image. Nonetheless, networks with higher sample classification accuracy will outperform less accurate sample classifiers when applied to full image classification problems.

We used the OSC network to identify marked samples, and the most common classification among those samples defined the otolith classification. In tests where unmarked otoliths were present, the samples selected by the OSC were passed to the binary network. If the average “marked” probability returned by the binary network across all selected samples for an image was less than 0.8, the OSC classification was overridden and the image was classified as unmarked. Figure 6 shows full image classification accuracy results using the OSC and AOC algorithms in scenarios with between two and five independent mark classes. The error bars the Clopper-Pearson interval for a binomial distribution given the number of images and successful classifications in cross validation [25].

**Figure 6.**
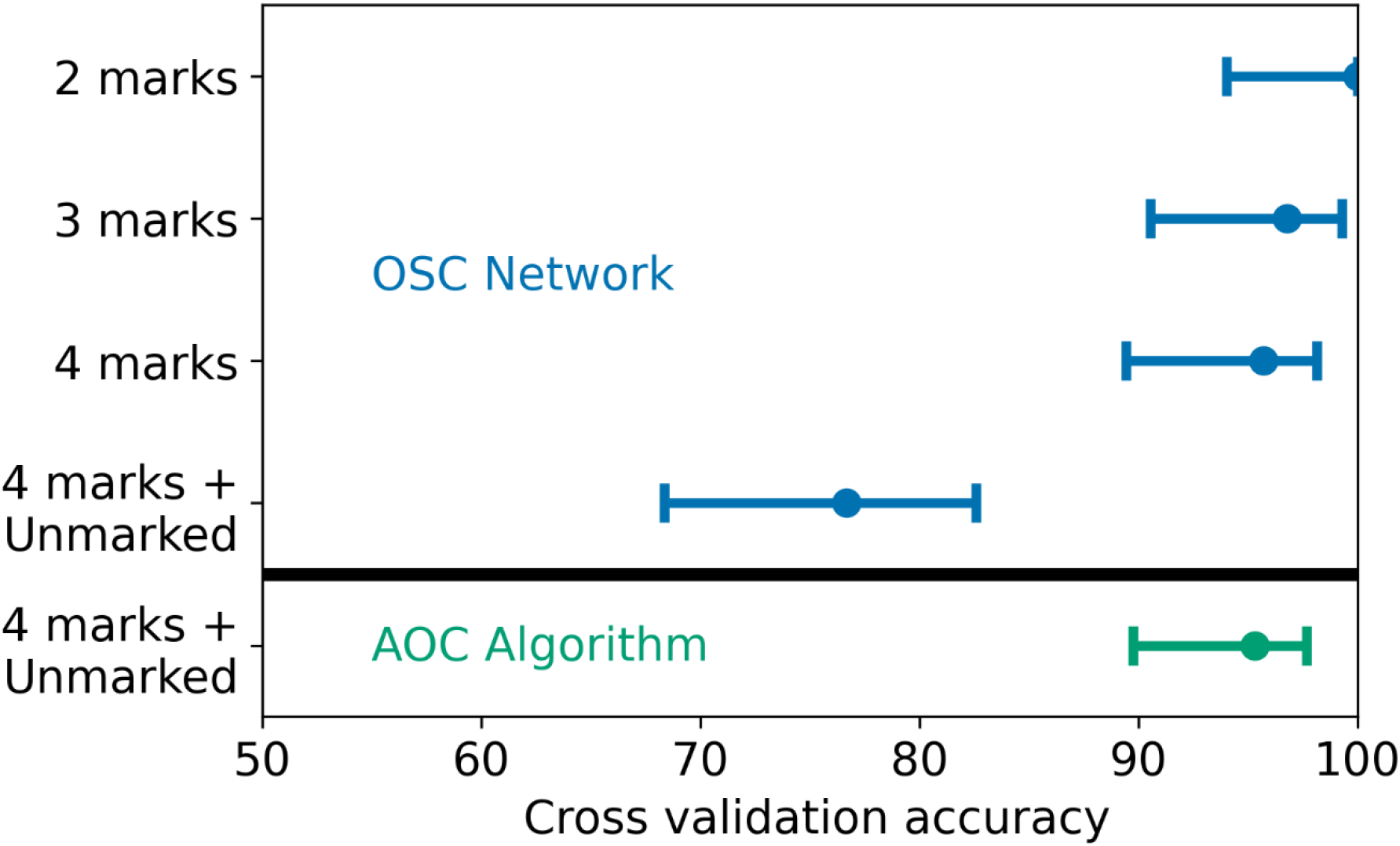
Full image classification accuracy of classification algorithms with different numbers of classes. The OSC network is used to select samples and classify images in the top four cases. The full AOC algorithm is implemented only in the final scenario.

The AOC algorithm is only necessary when the classes include unmarked otoliths. In scenarios without an unmarked class, the accuracy of image classification in the test set is above 95% using only the OSC. However, when unmarked images are included, none of those otoliths are classified correctly, resulting in an accuracy less than 80% (20% of the validation and test set images were unmarked). Adding the adversarial step to training allows unmarked images to be classified correctly at a similar rate to marked otoliths.

Table 1 summarizes the results of cross validation using the AOC algorithm in a confusion matrix. The row indicates the correct classification for each image, and the column indicates the classification chosen by the algorithm. In total 7 out of 150 images were classified incorrectly yielding an overall accuracy of 95%. The threshold used to separate unmarked images from marked images can be adjusted to reduce the probability of missing a marked image, or to maximize overall classification accuracy. In this case it was set to maximize classification accuracy in the cross validation data set, and the same setting was used for evaluating classification accuracy in the test set.

**Table 1.**
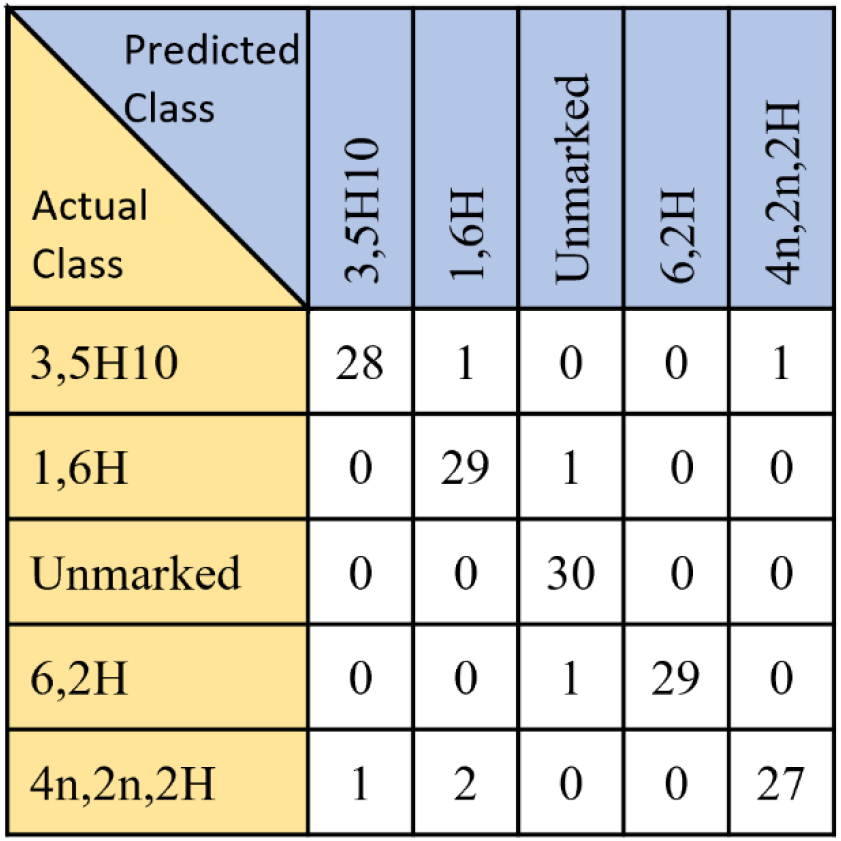
AOC cross validation results

| Predicted<br>Class<br>Actual<br>Class | 3,5H10 | 1,6H | Unmarked | 6,2H | 4n,2n,2H |
| --- | --- | --- | --- | --- | --- |
| 3,5H10 | 28 | 1 | 0 | 0 | 1 |
| 1,6H | 0 | 29 | 1 | 0 | 0 |
| Unmarked | 0 | 0 | 30 | 0 | 0 |
| 6,2H | 0 | 0 | 1 | 29 | 0 |
| 4n,2n,2H | 1 | 2 | 0 | 0 | 27 |

Table 2 shows the confusion matrix resulting from the same AOC algorithm applied to the test set. 7 out of 100 images were misclassified, resulting in an overall accuracy of 93%. Misclassification of marked images as unmarked persisted in the test set.

**Table 2.** AOC test set results

| Predicted<br>Class<br>Actual<br>Class | 3,5H10 | 1,6H | Unmarked | 6,2H | 4n,2n,2H |
| --- | --- | --- | --- | --- | --- |
| 3,5H10 | 16 | 0 | 2 | 2 | 0 |
| 1,6H | 0 | 19 | 1 | 0 | 0 |
| Unmarked | 0 | 1 | 19 | 0 | 0 |
| 6,2H | 0 | 0 | 0 | 19 | 1 |
| 4n,2n,2H | 0 | 0 | 0 | 0 | 20 |

## 4 Discussion

Automated otolith mark classification is challenging because the intra-class variability in ring location is comparable to the inter-class variability, and because the mark may only be clear in a small area of the full image. Figure 7.A and Figure 7.B show samples drawn from a 1,6H otolith, while Figure 7.C and Figure 7.D show samples from a 6,2H otolith. In Figure 7.A, the mark spans less than 400 of the 800 pixels along the long axis of the sample, while in Figure 7.B the mark spans approximately 600 of the 800 pixels. Furthermore, one of the rings in Figure 7.A is obscured by unintended variations in the otolith so that only five rings in the main band are easily identified. This sample could be misidentified as a 3,5H or 6,2H sample. Figure 7.C and Figure 7.D both show 6,2H marks, but the intensity and width of the rings vary between the two samples.

**Figure 7.**
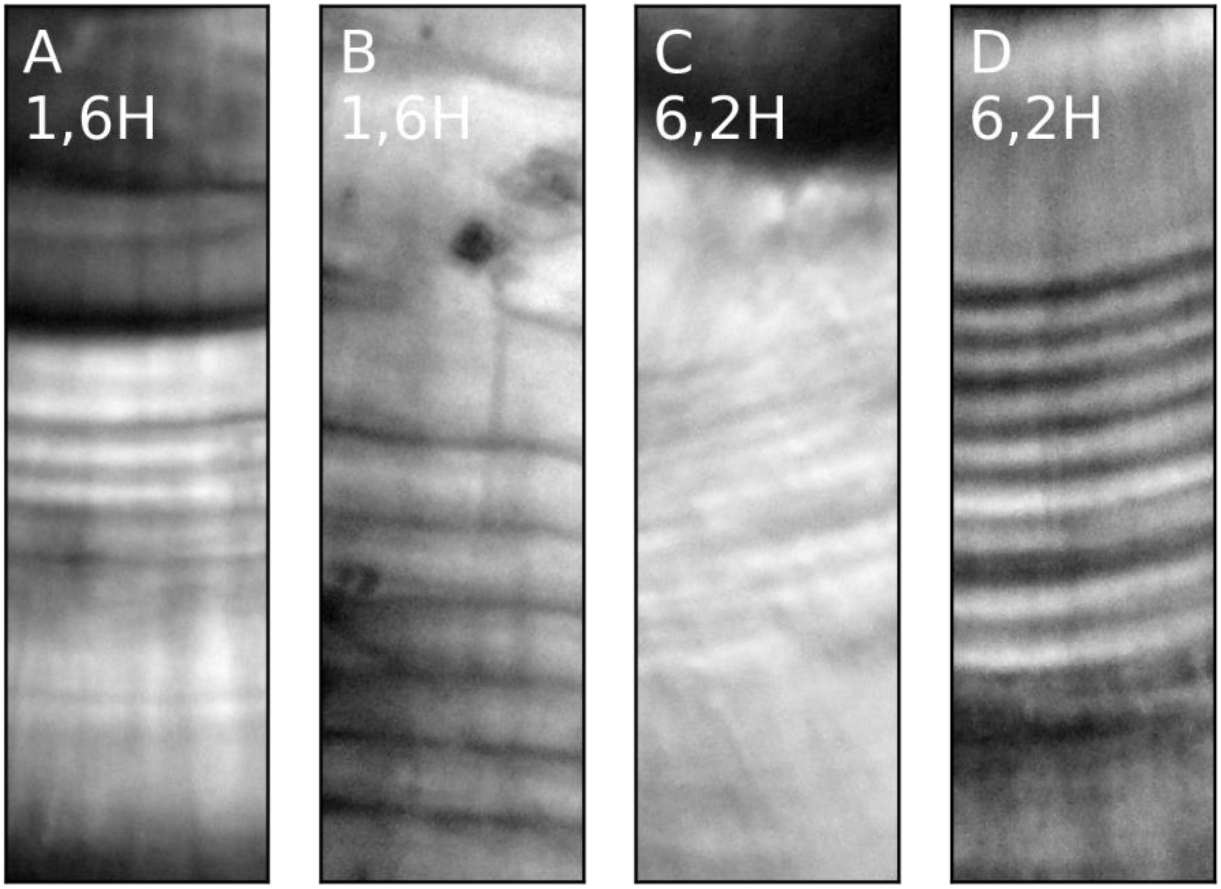
Examples of challenging samples extracted from otolith images. A and B show samples from a 1,6H otolith, while C and D show samples from a 6,2H otolith.

Early classification algorithms relied on a ring spacing threshold to distinguish between different marks [11]. The intraclass variability shown in Figure 7 makes that type of approach inadequate for these marks. The neural network approach supports higher level relationships between rings that are difficult to encode directly. The algorithm does not require a specific spacing between rings, nor does it require meeting an intensity or contrast threshold. The network incorporates data from throughout each sample to develop a classification guess for that sample without any explicit ring counting.

The AOC algorithm uses brute force to identify marked samples within an image: samples are iteratively drawn from the image at full resolution and considered independently. This approach is computationally intensive and might be improved in the future by using low resolution thresholds to quickly filter out unmarked areas of the image. The strength of the approach is that it reliably identifies marked samples throughout the sample that are all considered in the image classification; if some rings are obscured as in Figure 7.A, the image can still be classified correctly if the full mark is detected elsewhere.

### 4.1 Sample size limitations

The primary limitation to classification accuracy is overfitting. Figure 8 shows the training and validation accuracy of 10 cross validation folds over 10 epochs for the otolith sample classifier network. The training accuracy approaches 100% in this case, while the validation accuracy averages 82% after 10 folds. This indicates that further reducing the degrees of freedom in the model or increasing the number of training samples may improve the accuracy of the network. Small variations to the algorithm were explored such as adding dropout layers and max pooling layers, but they were found to have little affect on the accuracy of the algorithm.

**Figure 8.**
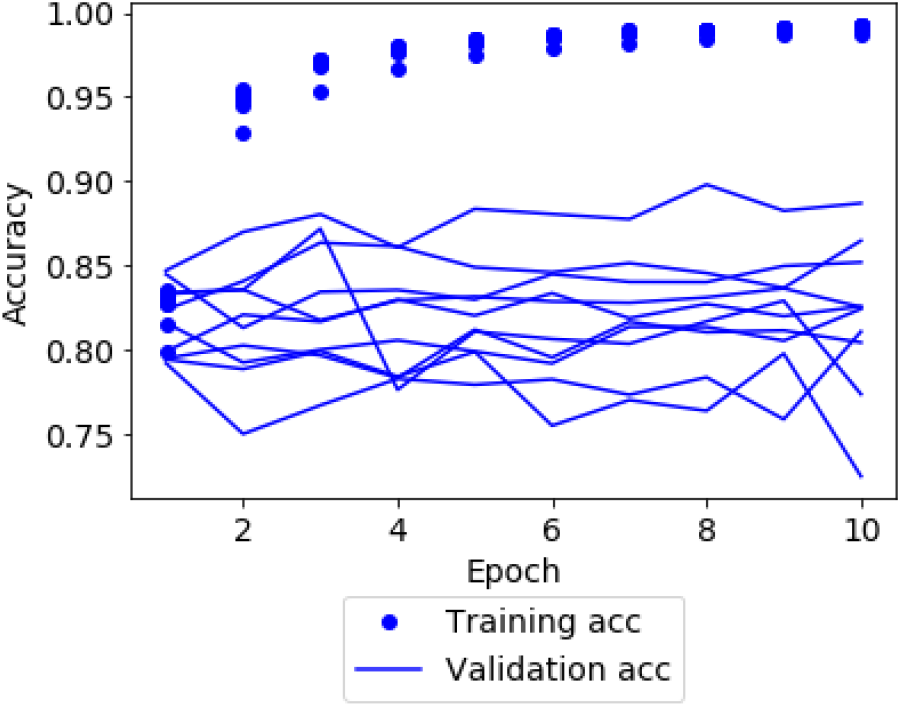
Training and validation accuracy for the OSC network with max pooling

In addition to limiting the accuracy of classifications, the small sample size limits the precision of accuracy estimates. With only 100 samples in the test set the accuracy was found to be 93%. We can conservatively estimate the uncertainty using the Clopper-Pearson interval for a binomial distribution [25]. The 95% confidence interval is 86-97% for the test set, and 90-98% for the cross validation results. This uncertainty in accuracy makes further improvements to the model difficult to assess with the existing sample size: a change to the model that halved the number of misclassified images would not result in a statistically significant improvement. Therefore, expanding the otolith image database will be critical to further improving the accuracy of the model.

### 4.2 Future work

Our work demonstrated the accuracy of the Adversarial Otolith Classifier with up to four unique marks mixed with unmarked otoliths. Future work should expand the training and test sets to include more marks to test the performance of the algorithm under the more complex environment observed in the field, where dozens of marks may be present in one fishery. The consistent accuracy of the method as the number of classes was increased from two to four marks suggests that the method may be effective at classifying more diverse marks, but further investigation will be necessary to demonstrate accuracy in more realistic conditions.

Our results also benefitted from using high quality images from otoliths of equal age for both the training and test sets. In practice otolith images used for classification by human readers often show the mark with less clarity. Furthermore, human readers refer to otolith images taken from salmon fry to classify otoliths retrieved from adult salmon. Future work should explore the accuracy of classification algorithms when using lower quality images from otoliths retrieved from salmon at different ages for training and testing.

While the present algorithm merely approaches the abilities of human readers, applying machine learning to a large set of otolith images could yield patterns that are not discernible to human readers. Identification of wild fish stocks based on otolith patterns may be possible, as well as signatures of traumatic events. These deeper applications that would provide insight beyond what can be achieved by human readers without computer assistance will be the next frontier in automated otolith reading.

## 5 Acknowledgements

The authors would like to thank Alan Murray, Whitney Chittenden, Tessa Frost and Matt Allen for contributing decades of otolith reading experience and procuring samples for this work.

## Supporting Information

**S1 File. Supporting Information.** Appendices to this manuscript.

**S1 Fig. Example of samples generated by the ImageDataGenerator (IDG).**

**S2 Fig. Overtraining of fine-tuned published networks.**

**S3 Fig. Overtraining of published transfer networks after 10 epochs without fine tuning.**

**S1 Table. Number of samples of each type included in database.**

**S2 Table. Layers in the classification network.**

**S3 Table. Layers in the final binary network.**

**S4 Table. Five-class accuracy of tested neural networks.**

**S5 Table. Fish brood years expected to return in 2020 by species.**

## Notes

https://doi.org/10.17882/84047

### Competing Interest Statement

The authors have declared no competing interest.

