## Supplemental Information for "Determining salmon provenance with automated otolith reading"

Kempy Energetics
306 Saint Anns Ave, Douglas, AK 99824

Otolith Marking and Reading Research
2847 Alaska Ave, Ketchikan, AK 99901

### Supplementary Information

3 Figures, 5 Tables

#### S1. Sample Preparation

Otolith samples were extracted from salmon caught by commercial fishers. Otoliths were cleaned with water then mounted on petrographic slides. The samples were then ground using a disk sander until the primordia of the otolith were visible when viewed at 200x magnification. The marks were then imaged through a microscope at 400x magnification.

#### S2. Algorithm development

The algorithms presented here were developed using a database of 250 otolith images. The database was used to iteratively train a classification algorithm, test the accuracy of the classifier using cross validation, and then update the training algorithm. The final accuracy of the algorithms was determined using a separate set of test images. These steps are listed explicitly below:

1. 50 otoliths from each of five classes were extracted, prepared and imaged with a magnification factor of 400.
2. 30 otoliths from each class were designated as the training and validation set, while the remaining 20 otoliths were designated as the test set.
3. Approximately 30 subsamples containing the mark were manually extracted from each otolith image in the training set. The samples were stored in a database.
4. A machine learning algorithm was trained using the database established in step 3.
5. The performance of the algorithm was estimated using cross validation
6. The algorithm accuracy was evaluated on the test set.

The subsequent sections present two families of algorithms that were developed by iterating between steps three and four. The first algorithm family used a transfer network built from published, deep neural networks trained with databases of millions of images. The accuracy of algorithms in this family proved to be limited but increasing the training set size may dramatically improve the methods' accuracy. Published transfer networks may also be more adaptable to new applications and otolith marks than the second algorithm family.

The second family of algorithms leveraged additional information about otolith structure to reduce the dimensionality of the training challenge and achieved more accurate results with the available training set. These methods reduced otolith mark samples to one dimension, and used a pair of networks to classify images. This family of algorithms will be referred to as 1D networks.

#### S2.1 Training database development

The algorithms require classifying the otolith images based on the mark without regard to the areas in the image that do not contain a mark. This motivated a training database that contained pre-cropped image samples centered on a visible hatchery mark. Approximately thirty such samples were extracted from each image in the training set.

A program was written to facilitate creating the database. The program displays images one at a time from a training directory. The user then clicks the location from which a sample should be extracted. The program uses a fourier transform algorithm to determine the rotation, centered on the point selected by the user, that results in the largest periodic variation. The program then draws a rectangle on the original image centered at the location clicked by the user and oriented to maximize the signal from the fourier transform algorithm. The user can then choose whether to store the sample to the database. The program is written in python and supported by a Jupyter notebook.

The final training database used in this analysis is described in Table S1. The number of unique samples selected in each image was determined by the area in which the mark was visible to the human eye. Samples were selected subjectively while maintaining a minimum 20-pixel separation from overlapping samples. Since the unmarked samples could be drawn from anywhere in the image, the samples were drawn randomly using an automated process rather than requiring human input. One hundred twenty unmarked samples from each unmarked image in the training set were included in the training database. Algorithms requiring an equal number of samples in each training class can down sample to the minimum number of samples in the database, or up-sample (through duplication or random generation) from the classes with less samples.

*Table S1 Number of samples of each type included in database*

| Mark type | Number of samples |
|---|---|
| 1,6H | 847 |
| 3,5H10 | 890 |
| 4n,2n,2H | 827 |
| 6,2H | 898 |
| Unmarked | 3600 |

While Table S1 shows over 800 samples in each class, it is important to note that the samples are not independent. All samples drawn from the same image are strongly correlated, with similar deviations from the ideal mark. While the additional samples are expected to improve the training in comparison to drawing just one sample from each image, they are not as valuable as samples drawn from distinct images.

Prior to developing the network algorithms defined below, numerous feature-based algorithms were developed that did not achieve the desired accuracy. For example, samples were averaged to one dimension, maxima in the second derivative were used to locate peak locations, and the distances between peaks were compared to the distances expected for each mark class. These efforts ultimately proved inadequate due to the complex variations between samples from the same image class. The distance between rings is not consistent between otoliths. Neither is the distance between rings within a single band: the first two rings might be separated by 20 pixels, while the second and third might be separated by 15 pixels. Yet, the marks are easily distinguished by eye.

The patterns that the eye catches are not related to the ring locations by a simple linear relationship. Rather, the pattern needs to be defined using a complex combination of features. In principle, those features could be defined directly and hard-coded into a classification algorithm. However, after investing significant effort, those features remained elusive in this work. Neural networks provided a method for using a computer to identify distinguishing features. In image classification, neural networks associate a weight with each pixel as well as groups of pixels and groups of groups of pixels. The computer randomly guesses each weight, checks the classification accuracy, then updates the weights. The process iterates for a predetermined number of "epochs." Once a neural network is well trained, the computer has a numerical representation of the relationship between pixel intensities and image classification.

Adding nodes and layers to a neural network allows for more complex patterns to be recognized resulting in more precise classification. However, additional nodes create degrees of freedom that require extra training data to constrain. Given the limited training data available here (~900 samples drawn from thirty images in each class), the neural network cannot have as many degrees of freedom as some of the networks trained on much larger datasets [1].

#### S2.2 Extracting marked samples from validation images

Otolith classification is a two step process. The first step is identifying sections of the image that contain a mark and extracting samples. The second step is classifying the image based on the samples that were selected. Accuracy can be measured at either step:

1. **Sample classification accuracy** is the fraction of selected samples that are classified correctly.
2. **Image classification accuracy** is the fraction of images classified correctly based on the classification scores for every selected sample from the image.

The following sections will refer to both types of accuracy in order to describe the performance of the algorithms.

#### S2.3 Transfer algorithms

The current state of the art image classification methods for novel classes with relatively small sample sizes use deep convolutional neural networks trained on a large database of unrelated images, with the top few layers re-trained or removed and replaced with new layers trained with the novel data set [2]. This approach is known as a "transfer network" because the knowledge stored in the original network is "transferred" to the final network in the fixed weights.

Many deep, pretrained networks (and their associated weights) are now available publicly. The Keras software package used throughout this work includes 10 such networks [3]. The pretrained networks are trained on large labeled image repositories. For example, the VGG19 network tested in the subsequent sections was trained with the ImageNet repository, which contains 1.3 million images divided into 1000 classes.

In addition to the pretrained networks, Keras provides tools for preprocessing images before submitting them to the network. These tools are leveraged in the following sections to explore the efficacy of using transfer networks with the database described in Section S2.1. The results of transferring three of those networks to otolith classification are examined in the following section.

##### S2.3.1 Preprocessing samples

The deep networks explored in this work require square inputs with up to 220-300 pixels on a side, depending on the specific network[1], significantly less than the 800x240 pixels stored for samples in the training database. Therefore, a copy of the database with all samples reduced to 200x60 pixels was established for use here.

The preprocessing algorithm uses the ImageDataGenerator (IDG) class from Keras. The quarter size sample training database was passed to the IDG. The IDG was used to randomly choose batches of samples from the database, rescale the pixel intensities to values between 0 and 1, and apply the following stochastic transformations:

- Rotations from 0° to 360°
- Horizontal shifts up to 20% of the sample width
- Vertical shifts up to 20% of the sample height
- Shear up to a factor of 0.2
- Zoom adjustments of up to 20%
- Vertical flips
- Horizontal flips

Pixels that were left undefined by the transformations (such as corner pixels after a rotation, or edge pixels after a translation) were filled with the nearest defined pixel intensity. Figure S1 provides three examples of samples generated by the IDG.

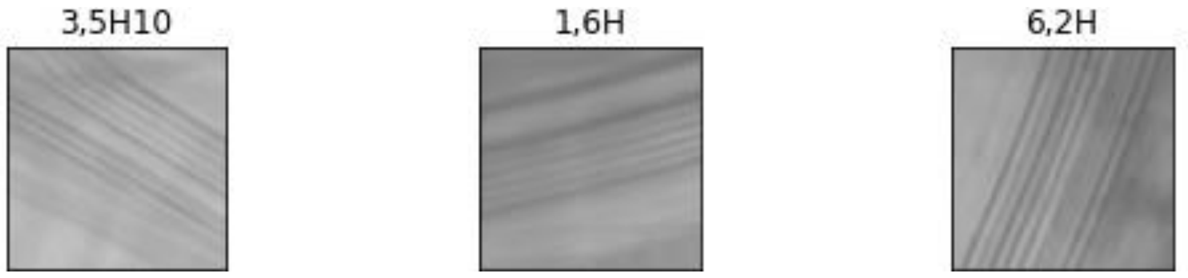


*Figure S1 Example of samples generated by the IDG*

The sample batches generated by the IDG were passed to the various transfer networks for training. Batch size was set to 20 samples, and 200 steps were specified per epoch. This results in exposing 4000 generated samples to the network per epoch (approximately equal to the number of unique samples in the training database).

---

[1] While the original networks were trained with a specific sized image, Keras uses the translation invariance of the networks to allow images up to the original training size.

##### S2.3.2 VGG19 transfer network

The VGG19 network was developed in 2014 and trained using the ImageNet database [1], [4]. The original VGG19 network consists of 16 convolutional layers and 3 fully connected layers. The network available in Keras contains all 16 convolutional layers, but the final fully connected layer is omitted. In this work, an additional dense layer with 256 nodes and relu activation was appended to the VGG19 network in addition to a 5 node dense soft max layer for the final classifications.

##### S2.3.3 InceptionV3 transfer network with fine tuning

The InceptionV3 network was developed to improve computational efficiency of image classification while supporting deeper networks [5]. The realization of the network supported by keras carries 314 layers (including layers with no parameters). The model was again initialized with weights from training with the ImageNet database.

Three additional layers were added to the InceptionV3 network for this work: a global average pooling layer, a dense layer with 1024 nodes and relu activation, and a dense 5 node layer with softmax activation. These new layers were trained for ten epochs with 200 steps per epoch using the sample generators defined in Section S2.3.1. After the new layers were trained, the top two training blocks of the original InceptionV3 network were also opened for training. The network was then trained for an additional 100 epochs.

##### S2.3.4 Xception transfer network with fine tuning

The Xception network has a similar architecture to the Inception network, but achieves 1% better validation accuracy using the ImageNet validation test set. The same transfer network architecture was used to implement Xception-based classification that was used with the Inception network. In addition, a second transfer network was invoked that used a dense layer with 128 nodes rather than 1024 nodes before the softmax layer.

#### S2.4 1D networks

The 1D networks described here use subject knowledge to dramatically reduce the dimensionality of the problem and eliminate the need to invoke public, pre-trained networks. Each 800x240 pixel sample stored in the database was reduced to a vector of 151 numbers. A compact, 2-layer convolutional neural network was then trained on the relatively low dimensional samples.

##### S2.4.1 1D network training

Each sample from the database of mark images described in Section S2.1 was augmented and adapted using the six steps described below.

1. Data augmentation: The training database was augmented by generating 20 variations of every sample by applying the following stochastic transformations:
    - rotations from -10 to +10 degrees
    - Height shifts of +/- 0.02
    - Vertical flips
2. De-trending and standard mean: The linear trend along the vertical axis was subtracted from each column of pixels

3. Normalized intensity variation: all residual pixel intensities were divided by the standard deviation of intensity across the full sample
4. 1-D samples: Every sample was averaged across the x-axis to produce a one dimensional sample
5. Smoothing and curvature emphasis: each 1-D sample was convolved with a 20 pixel smoothing kernel and a finite sample second derivative was applied to emphasize regions of the image with high curvature
6. Down sampling: Every fourth pixel was saved to a training set

The result of processing is an array of shape [20, 151] for each sample in the original training database. The first dimension (20) defines the number of synthetic samples derived from each original sample, and 151 is the length of the vector stored for each synthetic sample.

##### S2.4.2 1D-Internal Transfer Network Classification

Networks considered in this work that consistently discriminated between marked samples tended to mis-classify unmarked samples. In order to overcome this limitation, a two-network approach was adopted. The first network selected samples that appeared to contain a mark from each image and classified them. These samples were then passed to a second network that provided a binary, “marked” or “unmarked” result. This approach improved overall accuracy by maintaining the marked classifications from the 5-class network while identifying some unmarked images that the 5-class network had misclassified.

The method is referred to as an “internal transfer network” because the binary network was first trained on the same training set as the 5-class network, and then the final layers were replaced and retrained using samples labeled with a mark by the 5-class network.

Training the internal transfer network classification algorithm proceeded in six steps:

1. Initialize a 5-class network to distinguish between samples of 3,5H10, 1,6H, 4n,2n,2H, 6,2H and unmarked samples
2. Train the 5-class network on a set of labeled samples
3. Initialize a binary network to distinguish between marked and unmarked samples
4. Train the binary network on a set of labeled samples
5. Select samples from training images identified as marked by the classification network
6. Retrain the final two layers of the binary network using the samples selected in step 5

Classification using the internal transfer network algorithm consisted of

1. Selecting marked samples using the 5-class network
2. Designating images as **marked or unmarked** using the binary network and the samples selected in step 1
3. Classifying marked images as **1,6H; 3,5H10; 6,2H; or 4n,2n,2H** using the sample classification scores from step 1

*The 5-class network consisted of the layers described in*

Table S2. The impact of changing the kernel size, adding max-pooling layers and implementing different loss functions is discussed in Section S3.1.

*Table S2 Layers in the classification network*

| Layer type | Kernel Size | Activation | Number of nodes |
|---|---|---|---|
| 1D Convolution | 3 | relu | 256 |
| 1D Convolution | 9 | relu | 128 |
| Flatten | - | - | - |
| Dense | - | relu | 256 |
| Dense | - | softmax | 5 |

The network was trained using the samples processed as described in Section S2.4.1. The model used the 'adam' optimizer supported by TensorFlow with a 'sparse_categorical_crossentropy' loss function and 10 training epochs [6].

The binary network consisted of the same first four layers described in Table S2 for the classification network, but the final Dense layer was modified to have only two nodes. It was initially trained using the same training array, optimization and loss function as the classification network.

Finally, samples were extracted from every image in the training and validation set at intervals of 120 pixels. The classification network was then used to generate an estimated probability of each sample having no mark, or one of the four expected marks. The 0.1% of samples from each image with the lowest probability of being unmarked were then saved. The final layer of the original binary network was replaced by the two layers shown in Table S3. The new layers were then trained with the samples selected by the 5-class network.

*Table S3 Layers in the final binary network*

| Layer type | Kernel Size | Activation | Number of nodes |
|---|---|---|---|
| Dense | - | relu | 56 |
| Dense | | softmax | 2 |

Training the initial binary network on the full training set and then fine tuning the result using samples selected by the classification algorithm leverages the full breadth of the training set to develop features for the binary network, and then fine tunes that network to distinguish between the samples selected by the classification network. The approach is inspired by the adversarial network approach, in which one network is designed to trick a second network and the two compete against each other in training [7].

### S3. Classification accuracy

This section presents results from variations of the algorithms included in the main text.

#### S3.1 5-class sample classification accuracy

All of the image classification algorithms considered here begin with a neural network that can distinguish between five classes: 3,5H10; 1,6H, Unmarked; 4n,2n,2H and 6,2H. Table S1S4 summarizes the results from 9 neural networks that were exposed to training and validation data. The two 1D networks are variations of the network presented in Section S2.4. All other networks consist of published image classifier networks with two additional dense layers designed to customize the class predictions to otoliths. "Validation Accuracy" indicates the accuracy measured using a single cross validation fold, while "Cross Validation Accuracy" shows the average validation accuracy across ten folds. Cross validation was not completed for some of the published networks due to their lack of promise.

*Table S4 Five-class accuracy of tested neural networks*

| Model | Validation Accuracy | Cross Validation Accuracy |
|---|---|---|
| 1D 151 node 5-class net | 0.79 | 0.85 |
| 1D 151 node 5-class net with max pooling | 0.83 | 0.82 |
| VGG19, 10 epochs | 0.47 | - |
| VGG19, 100 epochs fine tuning | 0.39 | 0.45 |
| Xception, 10 epochs | 0.25 | - |
| Xception, 100 epochs fine tuning | 0.19 | 0.30 |
| Xception, 10 epochs with 128 dense nodes in top layer | 0.25 | - |
| Xception, 100 epochs with 128 dense nodes in top layer with fine tuning | 0.18 | - |
| InceptionV3, 10 epochs | 0.36 | - |
| InceptionV3, 100 epochs, fine tuning | 0.25 | 0.39 |

##### S3.1.1 Validation accuracy of published transfer networks

The VGG19, Xception and Inception networks were chosen due to their demonstrated performance across a broad range of image classes, their availability within the Keras package, and the variety of network structures they represent. **Error! Reference source not found.** shows that for all of the networks with fine-tuning, the training accuracy increased throughout 100 epochs of training, while the validation accuracy did not improve. This indicates overfitting: there are too many degrees of freedom to constrain with the available training data set. Figure S2.C shows results of running a fine tuning algorithm with a relatively small (128 node) dense layer added to the underlying Xception network. The network was still clearly overfit, suggesting that the limitation cannot be avoided by reducing the dimensionality of the transfer network.

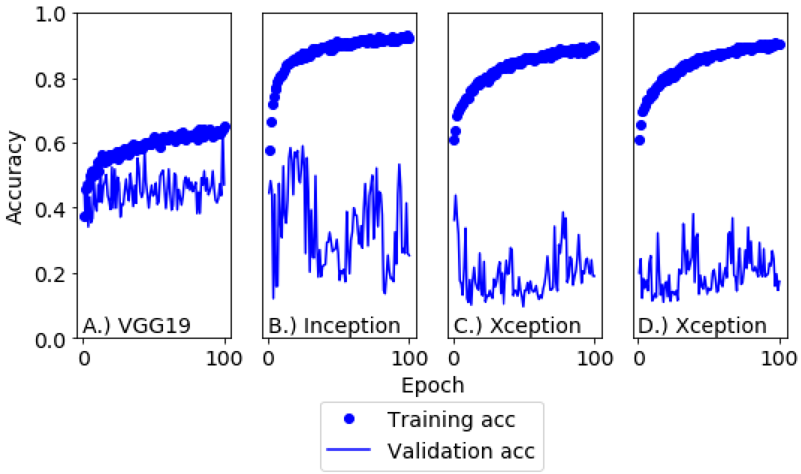


*Figure S2 Overtraining of fine-tuned published networks. A.) VGG19 model with a 256 node dense layer B.) Fine-tuned Inception model with 1024 node dense layer C.) Fine tuned Xception model with 1024 node dense layer D.) Fine tuned Xception model with 128 node dense layer*

Figure 3 shows that the overtraining is reduced when the networks are not fine tuned and the number of training epochs is reduced to 10. Nonetheless, the training accuracy increases throughout the 10 epochs in all four cases shown in Figure 3, while the validation accuracy remains constant or declines. The VGG19 network performs markedly better than the Inception and Xception networks (perhaps because the original network has less degrees of freedom and less depth than the Inception and Xception networks), but still performs much worse than the 1D 151 node networks.

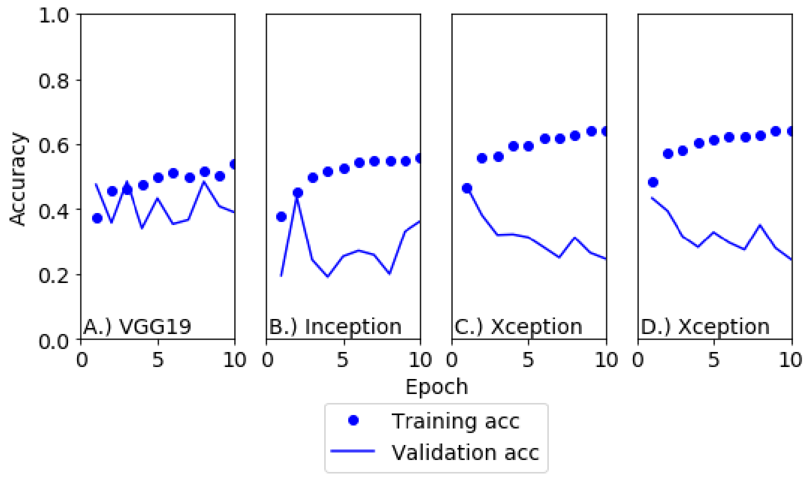


*Figure 3 Overtraining of published transfer networks after 10 epochs without fine tuning A.) VGG19 model with a 256 node dense layer B.) Inception model with a 1024 node dense layer C.) Xception model with 1024 node dense layer D.) Xception model with 128 node dense layer*

#### S4. Otolith marking locations

Figure 1 in the main text shows the number of unique marks expected to return to hatcheries throughout the Pacific Rim in 2020. The data were acquired from the North Pacific Anadromous Fish Commission (NPAFC) database of hatchery marks [8]. Marks were extracted from the database by species for the brood years listed in Table 5. The NPAFC records hatchery names but not locations, so the hatcheries were located by searching for their names using Google Maps. In some cases the hatchery itself could not be located, so the hatchery was placed in the watershed of the same name.

*Table 5 Fish brood years expected to return in 2020 by species.*

| Species | Brood years included |
|---|---|
| Coho | 2017 |
| Chinook | 2015-2017 |
| Chum | 2015-2017 |
| Sockeye | 2015-2017 |
| Masu | 2017-2018 |
| Pink | 2018 |
